# Body mass change over winter is consistently sex-specific across roe deer (*Capreolus capreolus*) populations

**DOI:** 10.1101/2022.09.09.507329

**Authors:** A.J.M. Hewison, N.C. Bonnot, J.M. Gaillard, P. Kjellander, J.F. Lemaitre, N. Morellet, M. Pellerin

## Abstract

In most polygynous vertebrates, males must allocate energy to growing secondary sexual characteristics, such as ornaments or weapons, that they require to attract and defend potential mates, impacting body condition and potentially entailing fitness costs.

We investigated sex differences in over winter body mass change across five intensively monitored populations of roe deer (*Capreolus capreolus*) with markedly contrasting environmental conditions. At winter onset, males weighed, on average, 8.4% (from 4.7% in the most northerly population to 11.6% in the most southerly one) more than females. However, across all populations, males fared worse over the winter than females, losing more (Sweden) or gaining less (France) mass, so that sexual mass dimorphism was virtually absent prior to the onset of spring.

Our findings reveal that the direction of over-winter change in mass of roe deer depends on winter severity, but that males are consistently more sensitive to this environmental constraint than females. As a result of this sex-specific change in body mass, sexual mass dimorphism is lowest at the onset of the territorial season. We suggest that allocation to antler growth and territory establishment drives this pattern, providing a likely explanation to account for the lower rates of male adult survival that are consistently reported in this weakly dimorphic species.

## Introduction

Although the difference in body mass of males and females is often considered as a fixed quantity for a given species, in reality, body mass of large herbivores varies seasonally in relation to resource abundance (Douhard et al. 2018) and the sex-specific schedule of allocation to reproduction (Apollonio et al. 2020). First, because of the greater energy requirements to sustain their larger body size, males are generally more susceptible to lose mass during periods of resource restriction, notably over winter in temperate areas (Clutton-Brock et al. 1982). Second, body condition is expected to fluctuate asynchronously between the sexes in relation to their different schedules of reproductive allocation (Stephens et al. 2009). In species that lack male parental care, females pay the costs of rearing offspring, notably during late gestation and early lactation, which generally coincide with the period of peak resource availability during late spring-early summer. Mothers can therefore offset this marked increase in energy expenditure by either increasing foraging activity (income breeder, sensu Jönsson 1997) or by mobilizing previously accumulated body reserves (capital breeder, sensu Jönsson 1997). In contrast, males must engage in contest competition to ensure access to mates, for example, through tending receptive females (Hogg 1984) or by defending a mating territory (Vanpé et al. 2009), and may lose substantial body condition as a result (Apollonio et al. 2020). In addition, to maximise their competitive ability, males must allocate substantial energy to developing secondary sexual traits including weapons such as antlers, which are regrown annually as a prerequisite to successful reproduction. Because allocation to these elements of male-male competition occurs earlier, typically prior to or during winter, when resources are less abundant in temperate areas, males are expected to adopt a capital breeder tactic (sensu Jönsson 1997), accumulating fat reserves during the season of highest resource abundance to offset the subsequent costs of reproduction (Williams et al. 2017).

The roe deer (*Capreolus capreolus*) is a weakly polygynous species (Vanpé et al. 2008) with a low level of sexual size dimorphism (Hewison et al. 2011) and relatively short antlers (Lemaitre et al. 2018), but where males are strongly territorial from late winter until the end of the summer (Vanpé et al. 2009). Unusually, roe deer males cast their antlers in late autumn which then regrow immediately over the following two to three months, so that the costs of allocation to antler growth are levied during the heart of winter. In contrast, roe deer females are not territorial, but are considered income breeders (Andersen et al. 2000), with very few fat reserves (Hewison et al. 1996), that breed every year irrespective of previous reproductive status (Andersen et al. 2000, Hewison and Gaillard 2001) and offset the annual costs of gestation and lactation during spring and summer through concurrent intake.

While we previously showed that immature juvenile roe deer of both sexes continued to gain mass at a similar rate over winter (Hewison et al. 2002 for two populations at 46-48°N latitude), no study has yet analyzed how sexual mass dimorphism of mature adults is impacted by winter harshness at a broad spatial scale. We addressed this knowledge gap by investigating how this unusual schedule of allocation to secondary sexual traits in males shapes sex differences in body mass change over the winter and, hence, the degree of sexual size dimorphism. We used body mass data derived from the intensive (> 7000 individuals), long-term (> 20 years) capture-mark-recapture monitoring of five roe deer populations living under markedly different ecological conditions to investigate the following predictions. First, because males must allocate to antler growth during the winter months, loss of body mass should be greater (or mass gain should be lower) in males than females so that sexual mass dimorphism is lowest at the onset of spring. Second, roe deer in the two Swedish populations should lose more body mass than those in the three French populations because of the much harsher winter in the north of the species’ range, although this may vary among years in relation to specific annual conditions.

## Materials & Methods

### Study sites

We focused on five intensively monitored populations of roe deer, three in France and two in Sweden, living on study sites with markedly different environmental conditions (Table 1). The two Swedish study sites are situated towards the northern limit of the species’ range, with harsh winter conditions, whereas the French study sites are within the southern part of the roe deer core range and have relatively mild winters. Otherwise, the study sites differ in terms of available habitat types, landscape structure and population density (Table 1).

**Table 1:** Study site characteristics of the roe deer populations. Sample size indicates the number of body mass measurements and the number of unique individual roe deer (i.e. the ratio indicates the mean number of measures per individual, see Table S10 for sample sizes per year). Julian date indicates when body mass was measured where 1 = Jan 1^st^ (see Bonnot et al. 2024 for data and code).

|  | Bogesund<br>(Sweden) | Grimsö<br>(Sweden) | Aurignac<br>(France) | Chizé<br>(France) | Trois-Fontaines<br>(France) |
| --- | --- | --- | --- | --- | --- |
| Latitude,<br>Longitude | 59°38'N,<br>18°28'E | 59°73'N,<br>15°47'E | 43°13'N,<br>0°52'E | 46°11'N,<br>0°34'W | 48°43'N,<br>4°55'E |
| Surface area (ha) | 2 600 | 8 000 | 7 500 | 2 614 | 1 360 |
| Habitat type | Mixed<br>agricultural | Boreal<br>coniferous forest | Mixed<br>agricultural | Deciduous<br>forest | Deciduous<br>forest |
| Snow cover (days) | 80 | 130 | 5 | <15 | <15 |
| January temperature (°C) | 3.7 | -1.3 | 4.9 | 5.6 | 3.1 |
| Years monitored | 1989-2016 | 1974-2017 | 2001-2023 | 1978-2015 | 1976-2015 |
| **Sample size:**<br>observations<br>(individuals) | 2432<br>(493) | 1516<br>(540) | 503<br>(361) | 5571<br>(3297) | 3887<br>(2564) |
| **Julian date:**<br>(start, end) | 2-92 | 1-99 | 5-74 | 4-84 | 4-73 |

### Body mass data

We collected data for all animals caught during routine capture-mark-recapture operations that took place each winter (January to March) over two to four decades depending on the study site (see Table 1). Animals were caught either in baited box traps (Sweden, see Kjellander et al. 2006 for more details) or drive nets (France, see Lemaître et al. 2018 and Hewison et al. 2009 for more details). They were subsequently manipulated by experienced handlers who recorded each individual’s sex, body mass (to the nearest 0.1 kg) and age (as either juveniles in their first winter i.e. around 8 months old, or adult i.e. older than 1.5 years old when both sexes have attained >90% of their asymptotic body mass, Hewison et al. 2011). Juveniles can be easily distinguished from older animals on the basis of the presence of a milk tooth at the third pre-molar (Ratcliffe & Mayle 1992). Animals were marked with individually numbered ear tags and, in some cases, collars, before being released on site.

### Ethical statement

All capture and marking procedures were done in accordance with local and European animal welfare laws. For Aurignac-VCG: prefectural order from the Toulouse Administrative Authority to capture and monitor wild roe deer and agreement no. A31113001 approved by the Departmental Authority of Population Protection. For Bogesund and Grimsö: the marking and handling of roe deer were approved by the Ethical Committee on Animal Experiments, Uppsala, Sweden (Current approval Dnr: C302/2012). For Chizé and Trois-Fontaines, the capture protocol for roe deer under the authority of the Office Français de la Biodiversité (OFB) was approved by the Director of Food, Agriculture and Forest (Prefectoral order 2009-14 from Paris). All procedures were approved by the Ethical Committee of Lyon 1 University (project DR2014-09, June 5, 2014).

### Data analysis

We analysed individual body mass of adult animals only in relation to sex and capture date defined as the number of days after 1^st^ January (hereafter, Julian date 1, see Bonnot et al. 2024). Although captures occasionally took place during October, November or December, we excluded these few data so as to consider a common winter start date across all five populations. However, because a given Julian date cannot be considered strictly equivalent between France and Sweden from a phenological point of view (e.g. different dates for spring vegetation green-up), we performed the analysis separately for each population. Hence, while the analysed range for Julian date started from 1 (i.e. January 1^st^), the end date differed somewhat among populations (see Table 1). Note that, as a consequence of this choice, it was not possible to formally test our second hypothesis with just five independent data points (populations).

Preliminary analysis indicated that body mass change over winter was adequately modelled as a linear function of date in all populations (little or no improvement in model fit when looking for non-linearity using quadratic, cubic or smoothing functions, see Table S6 in Appendix), and that including exact age did not influence the outcome (analyses restricted to known aged individuals, results not presented). Therefore, to evaluate sex-specific body mass trajectories over winter, we built linear mixed models with the lme4 (Bates et al. 2015) package in R where the full model contained sex, Julian date and their two-way interaction. We first scaled Julian date for each population by centering (i.e. subtracting each value from the mean Julian date) and then dividing it by its standard deviation. For the Aurignac-VCG population only, we also included the spatial sector of capture as a two-modality fixed factor (mixed vs. open habitat) to control for body mass differences in relation to landscape structure at this study site (i.e. roe deer heaviest in open areas, Hewison et al. 2009); note, we did not include animals caught in the strict forest sector because of systematic differences in capture date among sectors). Finally, we initially included individual identity (to control for repeated measures) and year (to control for annual variation in conditions) as random effects on both the intercept and the slope. While these models successfully converged in two out of five cases, the low number of repeated measures of individuals (Table 1) precluded convergence for the three French populations. Therefore, to investigate whether issues of pseudo-replication might affect model selection for these populations, we re-ran the analysis on a reduced data set that included a single observation per individual (with year as a random effect on both the intercept and slope). As we obtained equivalent results with this approach (same model selected, essentially identical parameter estimates), below we present the analysis based on the full data set in the main text, with the equivalent analysis on the reduced data set provided in the Appendix (Tables S7-9, Fig. S1). We performed model selection in relation to AIC values and weights for the candidate model set. For each population, we retained the model with the lowest AIC value as long as it differed by at least 2 points from any simpler competing model (see Arnold 2010).

## Results

In all five populations, the best supported model describing over-winter variation in body mass consistently included the sex by date interaction (for all five populations, ΔAIC > 3.5 compared to the second-best model), showing that average change in body mass over winter differed between males and females (see Tables S1-S5 for scaled parameter estimates). More specifically, in the two Swedish populations, body mass (mean ± sd) of males decreased by −21.1 g (± 3.1, Bogesund, and −21.5 g (± 3.7, Grimsö) per day between 1^st^ January and the end of the winter, while this decrease was much less marked for females (−4.5 ± 2.8 g and −12.0 ± 3.7 g /day, respectively). In contrast, in the three French populations, female body mass increased by between 14.2 g (± 3.1, Trois-Fontaines) and 25.8 g (± 6.1, Aurignac-VCG) per day over winter, while that of males remained more or less constant (from −2.3 ± 3.0 g/day at Chizé to 3.9 ± 4.3 g/day at Trois-Fontaines). As a result, while males were clearly heavier, on average, than females at the onset of winter in all five populations, albeit more pronouncedly in France (mean ± se: Chizé: 23.0 ± 0.2 kg for males vs. 20.7 ± 0.2 kg for females; Trois-Fontaines: 25.0 ± 0.2 kg for males vs. 22.8 ± 0.2 kg for females; Aurignac-VCG: 23.9 ± 0.3 kg for males vs. 21.4 ± 0.2 kg for females, i.e. a sexual mass dimorphism of about 10%) than in Sweden (Bogesund: 24.8 ± 0.2 kg for males vs. 23.5 ± 0.2 kg for females; Grimsö: 26.3 ± 0.2 kg for males vs. 25.1 ± 0.2 kg for females, i.e. a sexual mass dimorphism of about 5%), by mid-March (Julian date = 74), males did not weigh substantially more than females in all populations (Fig. 1). Finally, at Aurignac-VCG only, the best supported model included an additive effect of sector, indicating that deer weighed, on average, 0.81 kg (± 0.2) more in the open sector than those in the partially wooded sector.

**Fig. 1:**
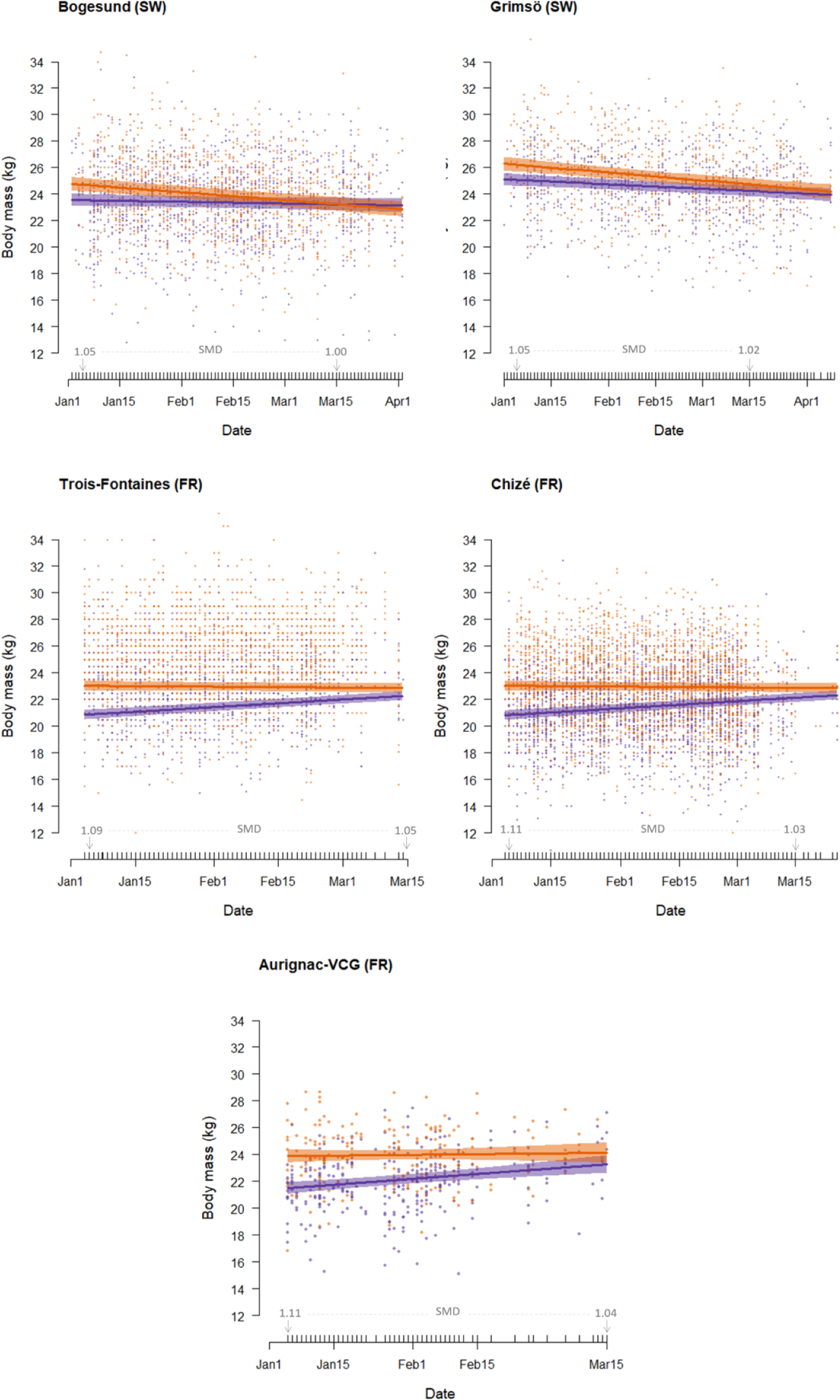
Body mass (kg) of male (orange) and female (purple) adults (>1.5 years old) in five intensively monitored populations of roe deer situated in Sweden (Bogesund, Grimsö) and France (Chizé, Trois-Fontaines, Aurignac-VCG) in relation to date over winter. Sexual mass dimorphism (SMD) calculated as the ratio of predicted male mass to female mass at the start (1^st^ January) and end (15^th^ March) of the winter period is indicated for each population.

Lastly, there was some evidence that over-winter body mass change varied among years to a greater extent in the Swedish populations compared to the French populations: the proportion of the variance attributable to the random effect of year on the slope of the body-mass-date relationship was 2.5-9.0 times higher in Bogesund and Grimsö than in the three French populations (Table 2).

**Table 2:** Variance attributed to the residual, fixed and random components of the model describing sex-specific over-winter variation in body mass across five populations of roe deer. To facilitate reliable comparison, we derived these variance components using an identical model structure for each population, with individual identity as a random effect on the intercept, and year as a random effect on both the intercept and the slope. The proportion of the total variance explained by the random effect of year on the slope is provided as a percentage.

|  | Residual | Fixed effects | Random effects |  |  |  |
| --- | --- | --- | --- | --- | --- | --- |
|  |  |  | *ID intercept* | *Year intercept* | *Year slope* | *% variance slope(Year)* |
| Bogesund (SW) | 1.67 | 0.18 | 5.97 | 0.63 | 0.05 | 0.53% |
| Grimsö (SW) | 1.83 | 0.30 | 4.92 | 0.60 | 0.06 | 0.72% |
| Trois-Fontaines (FR) | 2.52 | 0.88 | 4.20 | 1.05 | 0.01 | 0.09% |
| Chizé (FR) | 1.87 | 0.64 | 4.00 | 0.88 | 0.01 | 0.08% |
| Aurignac-VCG (FR) | 0.94 | 1.16 | 3.49 | 0.06 | 0.01 | 0.21% |

## Discussion

From the analysis of the body mass of over 7000 individual roe deer living along a gradient of ecological conditions in terms of winter harshness, from near the northern-most extreme to the southern part of their core geographical range, we found strong support for both our predictions, i/ that over-winter body mass change was sex-specific whereby males lost more (or gained less) than females, so that sexual dimorphism in mass was virtually absent by the end of the winter; ii/ but that this pattern was strongly modulated by winter severity such that while roe deer in Sweden lost mass, on average, those in France gained mass. There was also some indication that over-winter body mass change was more variable among years in the Swedish populations compared to the French populations, providing additional support for the latter hypothesis. The costs of allocation to sex-specific reproductive schedules likely drive seasonal variations in the degree of sexual mass dimorphism in this weakly polygynous ungulate.

### On the energetic cost of allocation to secondary sexual traits during winter

In polygynous mammals, reproductive effort during the mating season can lead to considerable loss of body mass in males (Apollonio et al. 2020; e.g. in red deer (*Cervus elaphus*): Yoccoz et al. 2002; in elephant seals (*Mirounga angustirostris*): Deutsch et al. 1990; in moose (*Alces alces*): Mysterud et al. 2005a). Indeed, reproductive males often abstain from feeding while they court and defend females or a mating territory (Mysterud et al. 2008). Similarly, although information on the costs of allocation to secondary sexual traits is sparse, antlers are smaller during less favourable years (Mysterud et al. 2005b), suggesting that growing these secondary sexual traits is costly. Here, we showed that male roe deer were consistently more constrained by winter resource restriction than females, losing around two to four times more mass in Sweden, while gaining up to six times less mass in France. As a result, by the onset of territorial season at the end of March (Hewison et al. 1998), sexual dimorphism in mass was virtually absent, with the average male only weighing about half a kilogram more than the average female across all five populations. While gestation in roe deer females begins in late December or early January following approximately 4.5 months of diapause (Aitken 1974), substantial allocation to foetal growth is concentrated in the latter third (April-May) so that fetuses weigh no more than a few grams during the winter period studied here (Beyes et al. 2017). We suggest that this over-winter decrease in sexual dimorphism of body mass is likely due to sex differences in the schedule of reproductive effort, in particular, the energetic costs to males of growing weapons and establishing a mating territory during the most resource-limited season (Williams et al. 2017).

### On the impact of winter severity for body mass change

While roe deer are consistently heavier in Sweden than France at winter onset (Fig. 1), the severity of conditions during the Scandinavian winter caused an average body mass loss of between 0.4 kg (females at Bogesund) and 2.1 kg (males at Grimsö). Note that these figures are likely conservative, as resource scarcity during winter may begin well before the New Year in northern environments depending on the annual timing of first snowfall. For an animal of around 20-25 kg this loss is clearly considerable and indicates that the capacity to store fat reserves and, therefore, seasonal fluctuations in body mass, are much greater in the north of its range than previously documented for this medium-sized income breeder (Kjellander et al. 2006). Indeed, there was also some indication in our data that over-winter body mass change varied from year to year somewhat more markedly in Sweden compared to France (see Table 2), presumably in response to the harshness of winter conditions in a given year. This is likely an adaptation to buffer against severe winters, as further south, in the heart of its range, over-winter body mass is generally stable and may even increase slightly (Hewison et al. 1996, 2002). Larger body size (Linstedt & Boyce 1985) and the capacity to store fat (Trondrud et al. 2021, Denryter et al. 2022) have been widely interpreted as adaptations which increase fasting endurance in response to the dramatic fluctuations of resource availability in highly seasonal environments. Although differences in the operational sex ratio across populations could theoretically influence relative priority of allocation to sexual secondary characters, such as antlers, in polygynous systems driven by variation in the intensity of male-male competition (Weir et al. 2011), this is highly unlikely in our specific case. Indeed, the roe deer is only weakly polygynous (Vanpé et al. 2008) and the adult sex ratio (number of males/total number of males and females) is ostensibly the same across the five populations (Chizé: 0.44; Trois-Fontaines: 0.47; Aurignac-VCG: 0.41; Bogesund: 0.43; Grimsö: 0.41). Larger body size has often been reported at higher latitudes within species of mammals (Ashton et al. 2000), in line with Bergmann’s rule, and is thought to reflect natural selection for greater thermoregulatory buffering in endotherms (He et al. 2023). Our data are also in line with this general pattern, but indicate that sexual selection is the ultimate driver of between-sex differences in over-winter body mass change, suggesting similar priority of energy allocation to this secondary sexual trait across hugely contrasted environments.

### On the life history implications of annual body mass loss during winter

The repeated annual cycles of fat accumulation and depletion that underpin a capital breeding tactic are predicted to carry life history costs (Houston et al. 2006). While there is clear evidence to indicate that roe deer females adopt an income breeder tactic relative to other large herbivores (Andersen et al. 2000), our results imply that males must accumulate body condition prior to winter to offset the energetic costs of antler growth and subsequent territory establishment and, in this sense, can be considered capital breeders relative to females (Apollonio et al. 2020). In polygynous mammals, allocation to traits that confer an advantage in contest competition for females are predicted to impose costs in terms of survival (Clinton & Leboeuf 1993). Previous work has established that, despite the low level of polygyny in roe deer (Vanpé et al. 2008), the sex difference in annual survival of adults is equivalent to that of more polygynous and size dimorphic ungulates (Gaillard et al. 1993). We suggest that the repeated energetic cost of allocating to secondary sexual traits every winter is a proximal driver that, together with the direct costs of territorial defense and male-male competition for mates, contributes to the survival deficit for males in this weakly dimorphic ungulate. Most deer species cast and re-grow antlers during spring, when resources are plentiful (Mysterud et al. 2005b). However, because of their unusual schedule of allocation to reproduction, roe deer males are repeatedly faced with a trade-off between maintaining accumulated mass to offset the costs of establishing and defending a mating territory in spring, a full four months prior to the rut, and growing antlers during the winter season of food scarcity. The relative importance of antler size, body mass and territory quality for determining male reproductive success has yet to be established. Despite the huge among individual variation in body mass at the onset of winter within all five populations (Fig. 1, Table 2), there was little evidence for individual variation in over-winter body mass change, although we could not formally evaluate this due to the very low number of repeat measures. Future investigations of inter-individual variation in over-winter body mass change in relation to environmental severity would be highly informative for understanding individual tactics of energy allocation to secondary sexual traits and their life history consequences.

## Acknowledgements

We thank the local hunting associations, the Fédération Départementale des Chasseurs de la Haute Garonne, and numerous co-workers and volunteers for their assistance. We also thank Denis Réale, Patrick Bergeron, Phil McLoughlin and Achaz von Hardenberg for constructive comments on previous versions of this manuscript. Author contributions: A.J.M.H., N.C.B., J.M.G. & P.K. conceived the ideas, designed the study and the methodology; all authors collected the data and discussed the analytical approach; N.C.B. analysed the data.; A.J.M.H. wrote the first draft of the manuscript and all authors contributed critically to revision.

## Funding

This work was supported by an August Larsson Visiting Fellowship to A.J.M.H. and the “DivinT” and “EVORA” projects funded by the Agence Nationale de la Recherche grants ANR-22-CE02-0020-03 and ANR-22-CE02-0021-03 to J.M.G, J.F.L., M.P., N.M. and A.J.M.H.

## Conflict of interest disclosure

The authors declare they have no conflict of interest relating to the content of this article. A.J.M.H. and N.C.B. are recommenders for PCIEcology.

## Data, scripts, code and supplementary information availability

Data and code can be found at Bonnot et al. (2024), while supplementary information is given in an Appendix at the end of this article.

## Appendix

### 1.1 Model selection

**Table S1a:** Model fit and selection (fixed and random effects, difference in AIC score compared to the best model, AIC weight) describing sex-specific over-winter variation in body mass in the **Bogesund** population. The selected model is shaded grey.

| Fixed effects | Random intercept | Random slope | ΔAIC | weight |
| --- | --- | --- | --- | --- |
| Sex + Julian date + Sex:Julian date | ID & Year | ID & Year | 0.0 | 1.00 |
| Sex + Julian date | ID & Year | ID & Year | 26.2 | 0 |
| Julian date | ID & Year | ID & Year | 27.2 | 0 |
| Sex | ID & Year | ID & Year | 37.7 | 0 |
| null | ID & Year | ID & Year | 38.4 | 0 |

**Table S1b:** Model summary for the selected model: parameter coefficients for the fixed effects (mean and standard error, s.e.) and variance components for the fixed, random and residual effects, describing sex-specific over-winter variation in body mass in the **Bogesund** population. Note that Julian date was centered and scaled (see text). The reference category is female.

| Parameter | Coefficient | s.e. | Variance |
| --- | --- | --- | --- |
| **Fixed effects** |  |  | **0.17** |
| Intercept | 23.38 | 0.22 |  |
| Sex (male) | 0.55 | 0.24 |  |
| Julian date | -0.10 | 0.06 |  |
| Sex (male):Julian date | -0.37 | 0.07 |  |
| **Random effects** |  |  | **6.65** |
| Individual identity (intercept) |  |  | 5.92 |
| Individual identity (slope) |  |  | 0.05 |
| Year (intercept) |  |  | 0.63 |
| Year (slope) |  |  | 0.04 |
| **Residual** |  |  | **1.63** |

**Table S2a:** Model fit and selection (fixed and random effects, difference in AIC score compared to the best model, AIC weight) describing sex-specific over-winter variation in body mass in the **Grimsö** population. The selected model is shaded grey.

| Fixed effects | Random intercept | Random slope | ΔAIC | weight |
| --- | --- | --- | --- | --- |
| Sex + Julian date + Sex:Julian date | ID & Year | ID & Year | 0.0 | 0.90 |
| Sex + Julian date | ID & Year | ID & Year | 4.4 | 0.01 |
| Julian date | ID & Year | ID & Year | 15.1 | 0.00 |
| Sex | ID & Year | ID & Year | 27.4 | 0.00 |
| null | ID & Year | ID & Year | 37.3 | 0.00 |

**Table S2b:** Model summary for the selected model: parameter coefficients for the fixed effects (mean and standard error, s.e.) and variance components for the fixed, random and residual effects, describing sex-specific over-winter variation in body mass in the **Grimsö** population. Note that Julian date was centered and scaled (see text). The reference category is female.

| Parameter | Coefficient | s.e. | Variance |
| --- | --- | --- | --- |
| **Fixed effects** |  |  | **0.30** |
| Intercept | 24.56 | 0.19 |  |
| Sex (male) | 0.74 | 0.22 |  |
| Julian date | -0.30 | 0.08 |  |
| Sex (male):Julian date | -0.24 | 0.09 |  |
| **Random effects** |  |  | **5.65** |
| Individual identity (intercept) |  |  | 4.92 |
| Individual identity (slope) |  |  | 0.06 |
| Year (intercept) |  |  | 0.60 |
| Year (slope) |  |  | 0.06 |
| **Residual** |  |  | **1.76** |

**Table S3a:** Model fit and selection (fixed and random effects, difference in AIC score compared to the best model, AIC weight,) describing sex-specific over-winter variation in body mass in the **Aurignac-VCG** population. The selected model is shaded grey.

| Fixed effects | Random intercept | Random slope | ΔAIC | weight |
| --- | --- | --- | --- | --- |
| Sex + Julian date + Sex:Julian date + Sector | ID & Year | Year | 0.0 | 0.87 |
| Sex + Julian date + Sector | ID & Year | Year | 4.0 | 0.12 |
| Sex + Sector | ID & Year | Year | 9.9 | 0.01 |
| Sex + Julian date + Sex:Julian date | ID & Year | Year | 12.0 | 0.00 |
| Sex + Julian date | ID & Year | Year | 15.3 | 0.00 |
| Sex | ID & Year | Year | 22.4 | 0.00 |
| Julian date + Sector | ID & Year | Year | 67.1 | 0.00 |
| Sector | ID & Year | Year | 73.8 | 0.00 |
| Julian date | ID & Year | Year | 82.2 | 0.00 |
| null | ID & Year | Year | 88.7 | 0.00 |

**Table S3b:** Model summary for the selected model: parameter coefficients for the fixed effects (mean and standard error, s.e.) and variance components for the fixed, random and residual effects, describing sex-specific over-winter variation in body mass in the **Aurignac-VCG** population. Note that Julian date was centered and scaled (see text). The reference category is female and the mixed sector.

| Parameter | Coefficient | s.e. | Variance |
| --- | --- | --- | --- |
| **Fixed effects** |  |  | **1.16** |
| Intercept | 21.30 | 0.20 |  |
| Sex (male) | 1.87 | 0.22 |  |
| Julian date | 0.43 | 0.10 |  |
| Sector (open) | 0.81 | 0.21 |  |
| Sex (male):Julian date | -0.37 | 0.15 |  |
| **Random effects** |  |  | **3.56** |
| Individual identity (intercept) |  |  | 3.49 |
| Year (intercept) |  |  | 0.06 |
| Year (slope) |  |  | 0.01 |
| **Residual** |  |  | **0.94** |

**Table S4a:** Model fit and selection (fixed and random effects, difference in AIC score compared to the best model, AIC weight,) describing sex-specific over-winter variation in body mass in the **Chizé** population. The selected model is shaded grey.

| Fixed effects | Random intercept | Random slope | ΔAIC | weight |
| --- | --- | --- | --- | --- |
| Sex + Julian date + Sex:Julian date | ID & Year | Year | 0.0 | 1.0 |
| Sex + Julian date | ID & Year | Year | 47.2 | 0.0 |
| Sex | ID & Year | Year | 66.2 | 0.0 |
| Julian date | ID & Year | Year | 375.9 | 0.0 |
| null | ID & Year | Year | 396.3 | 0.0 |

**Table S4b:** Model summary for the selected model: parameter coefficients for the fixed effects (mean and standard error, s.e.) and variance components for the fixed, random and residual effects, describing sex-specific over-winter variation in body mass in the **Chizé** population. Note that Julian date was centered and scaled (see text). The reference category is female.

| Parameter | Coefficient | s.e. | Variance |
| --- | --- | --- | --- |
| **Fixed effects** |  |  | **0.64** |
| Intercept | 21.44 | 0.16 |  |
| Sex (male) | 1.52 | 0.08 |  |
| Julian date | 0.33 | 0.04 |  |
| Sex (male):Julian date | -0.37 | 0.05 |  |
| **Random effects** |  |  | **4.88** |
| Individual identity (intercept) |  |  | 4.00 |
| Year (intercept) |  |  | 0.88 |
| Year (slope) |  |  | 0.01 |
| **Residual** |  |  | **1.87** |

**Table S5a:** Model fit and selection (fixed and random effects, difference in AIC score compared to the best model, AIC weight,) describing sex-specific over-winter variation in body mass in the **Trois-Fontaines** population. The selected model is shaded grey.

| Fixed effects | Random intercept | Random slope | ΔAIC | weight |
| --- | --- | --- | --- | --- |
| Sex + Julian date + Sex:Julian date | ID & Year | Year | 0.0 | 0.86 |
| Sex + Julian date | ID & Year | Year | 3.7 | 0.14 |
| Sex | ID & Year | Year | 14.1 | 0.00 |
| Julian date | ID & Year | Year | 322.1 | 0.00 |
| null | ID & Year | Year | 333.0 | 0.00 |

**Table S5b:**
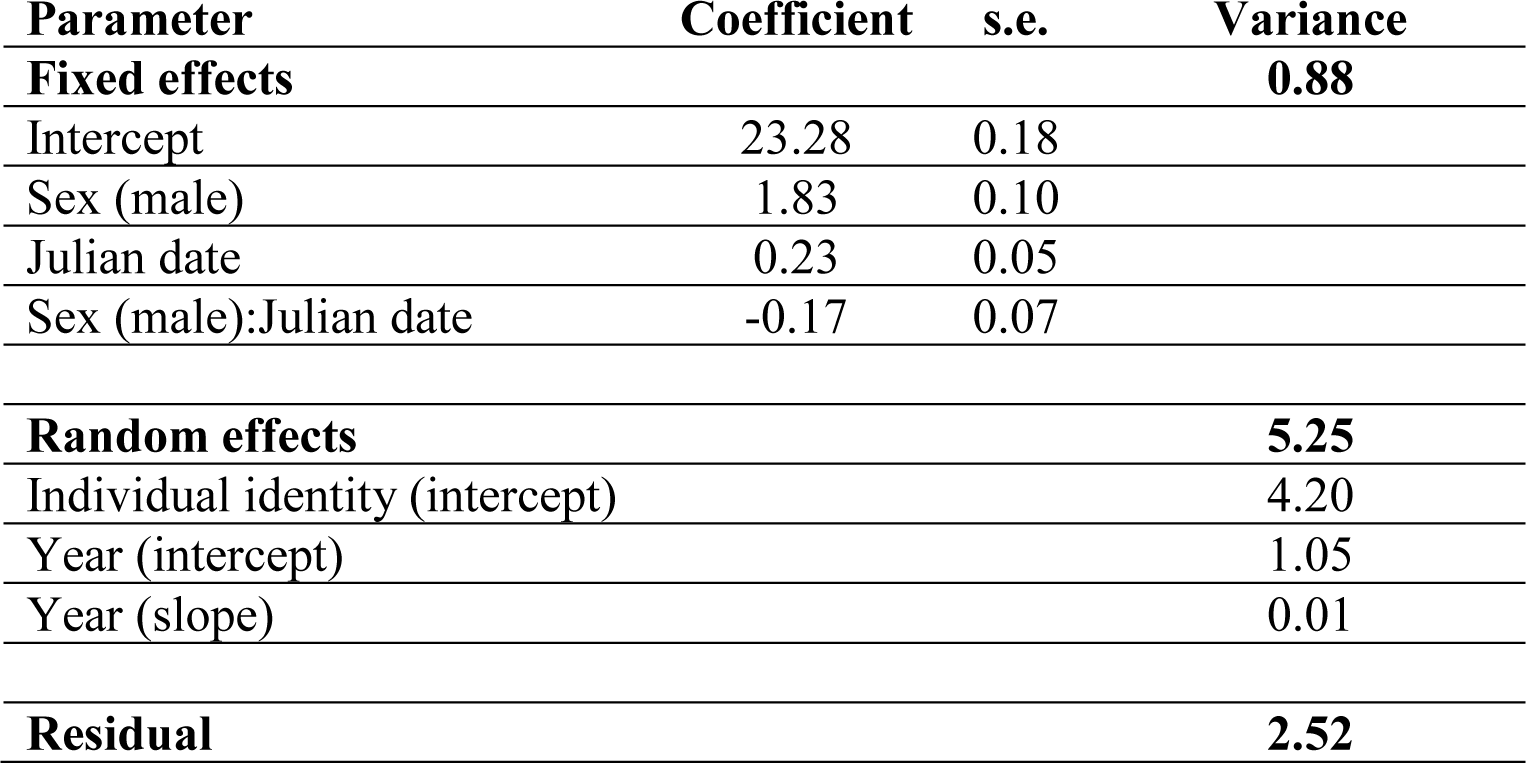
Model summary for the selected model: parameter coefficients for the fixed effects (mean and standard error, s.e.) and variance components for the fixed, random and residual effects, describing sex-specific over-winter variation in body mass in the **Trois-Fontaines** population. Note that Julian date was centered and scaled (see text). The reference category is female.

### 1.2 Test for non-linear variation in over-winter body mass

In preliminary analysis, in addition to a linear function, we modelled body mass change over winter with quadratic, cubic or smoothing functions of date using the ‘lme4’ and ‘gamm4’ packages (Bates et al. 2015; Wood & Scheipl 2020) in R. All models included sex, Julian date and their two-way interaction, as well as individual identity (to control for repeated measures on individuals) and year (to control for annual variation in conditions) as random factors. In three out of five cases, the linear function provided the best fit (Table S6), whereas the cubic model provided a somewhat better fit in the Grimsö population, while the quadratic model performed slightly better in the Aurignac-VCG population. Because this improvement was marginal from a biological point of view, and to facilitate comparison among populations, we present results from linear models in the main text, i.e. assuming that the rate of change in body mass during winter is constant over the entire study window.

Wood, S. & Scheipl, F. (2020) gamm4: Generalized Additive Mixed Models using ‘mgcv’ and ‘lme4’. R package version 0.2–6.

Bates, D., Maechler, M., Bolker, B. & Walker, S. (2015) Fitting linear mixed-effects models using lme4. Journal of Statistical Software, 67, 1–48.

**Table S6:** Comparison of model fit (AIC values) for the best supported model describing sex-specific over-winter variation in body mass in five roe deer populations (i.e. mass ∼ sex * Julian date, with an additive effect of sector for the Aurignac-VCG population only, see main text) when the relationship between body mass and date was modelled as either a linear effect, a quadratic effect, a cubic effect, or as a smoothing spline in a general additive mixed model framework). The selected model is indicated in bold.

| Population | linear | quadratic | cubic | GAMM |
| --- | --- | --- | --- | --- |
| Bogesund | **9526.9** | 9553.2 | 9530.4 | 9543.0 |
| Grimsö | 6348.3 | 6348.6 | **6342.7** | 6364.8 |
| Aurignac-VCG | 2057.3 | **2056.3** | 2058.6 | 2130.5 |
| Chizé | **24095.6** | 24146.1 | 24102.4 | 24439.6 |
| Trois-Fontaines | **17777.2** | 17782.8 | 17783.7 | 18109.9 |

### 2.1 Model selection on a reduced data set, featuring a single observation per individual (with year as a random effect on both the intercept and slope) for the three populations where the full model (which also included a random effect of individual identity on both the intercept and slope) did not converge

**Table S7:** Model fit and selection (fixed and random effects, difference in AIC score compared to the best model, AIC weight) on the reduced data set, describing sex-specific over-winter variation in body mass in the **Aurignac-VCG population**. The selected model is shaded grey.

| Fixed effects | Random intercept & slope | ΔAIC | weight |
| --- | --- | --- | --- |
| Sex + Julian date + Sex:Julian date + Sector | Year | 0.0 | 0.63 |
| Sex + Julian date + Sector | Year | 2.0 | 0.23 |
| Sex + Sector | Year | 3.1 | 0.13 |
| Sex + Julian date + Sex:Julian date | Year | 9.6 | 0.01 |
| Sex + Julian date | Year | 11.1 | 0.00 |
| Sex | Year | 14.7 | 0.00 |
| Julian date + Sector | Year | 64.8 | 0.00 |
| Sector | Year | 65.5 | 0.00 |
| Julian date | Year | 79.0 | 0.00 |
| null | Year | 82.3 | 0.00 |

**Table S8:** Model fit and selection (fixed and random effects, difference in AIC score compared to the best model, AIC weight) on the reduced data set, describing sex-specific over-winter variation in body mass in the **Chizé** population. The selected model is shaded grey.

| Fixed effects | Random intercept & slope | ΔAIC | weight |
| --- | --- | --- | --- |
| Sex + Julian date + Sex:Julian date | Year | 0.0 | 1.0 |
| Sex + Julian date | Year | 26.7 | 0.0 |
| Sex | Year | 31.2 | 0.0 |
| Julian date | Year | 329.9 | 0.0 |
| null | Year | 335.9 | 0.0 |

**Table S9:** Model fit and selection (fixed and random effects, difference in AIC score compared to the best model, AIC weight) on the reduced data set, describing sex-specific over-winter variation in body mass in the **Trois-Fontaines** population. The selected model is shaded grey.

| Fixed effects | Random intercept & slope | ΔAIC | weight |
| --- | --- | --- | --- |
| Sex + Julian date + Sex:Julian date | Year | 0.0 | 0.50 |
| Sex + Julian date | Year | 0.3 | 0.42 |
| Sex | Year | 3.7 | 0.08 |
| Julian date | Year | 291.9 | 0.00 |
| null | Year | 298.3 | 0.00 |

### 2.2 Estimated over-winter changes in body mass in the three French populations (Chizé, Trois-Fontaines, Aurignac-VCG) based on a reduced data set, featuring a single observation per individual

**Fig. S1:**
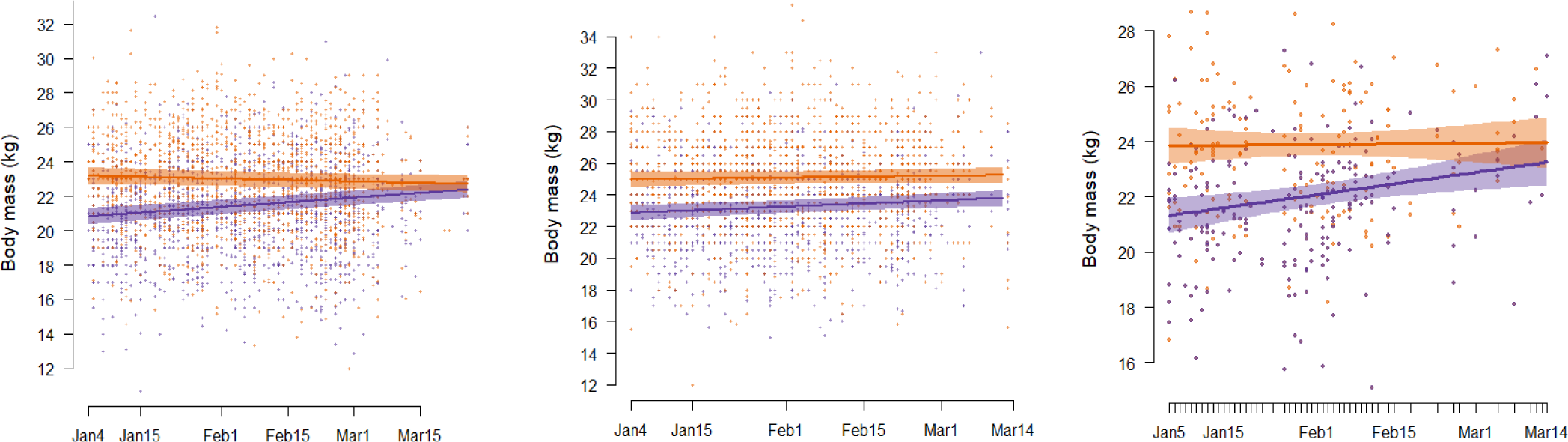
Body mass (kg) of male (purple) and female (orange) adults (>1.5 years old) in three intensively monitored populations of roe deer situated in France (Chizé, Trois-Fontaines, Aurignac-VCG) in relation to date over winter (based on the model selection presented in Tables S7-S9).

**Table S10:**
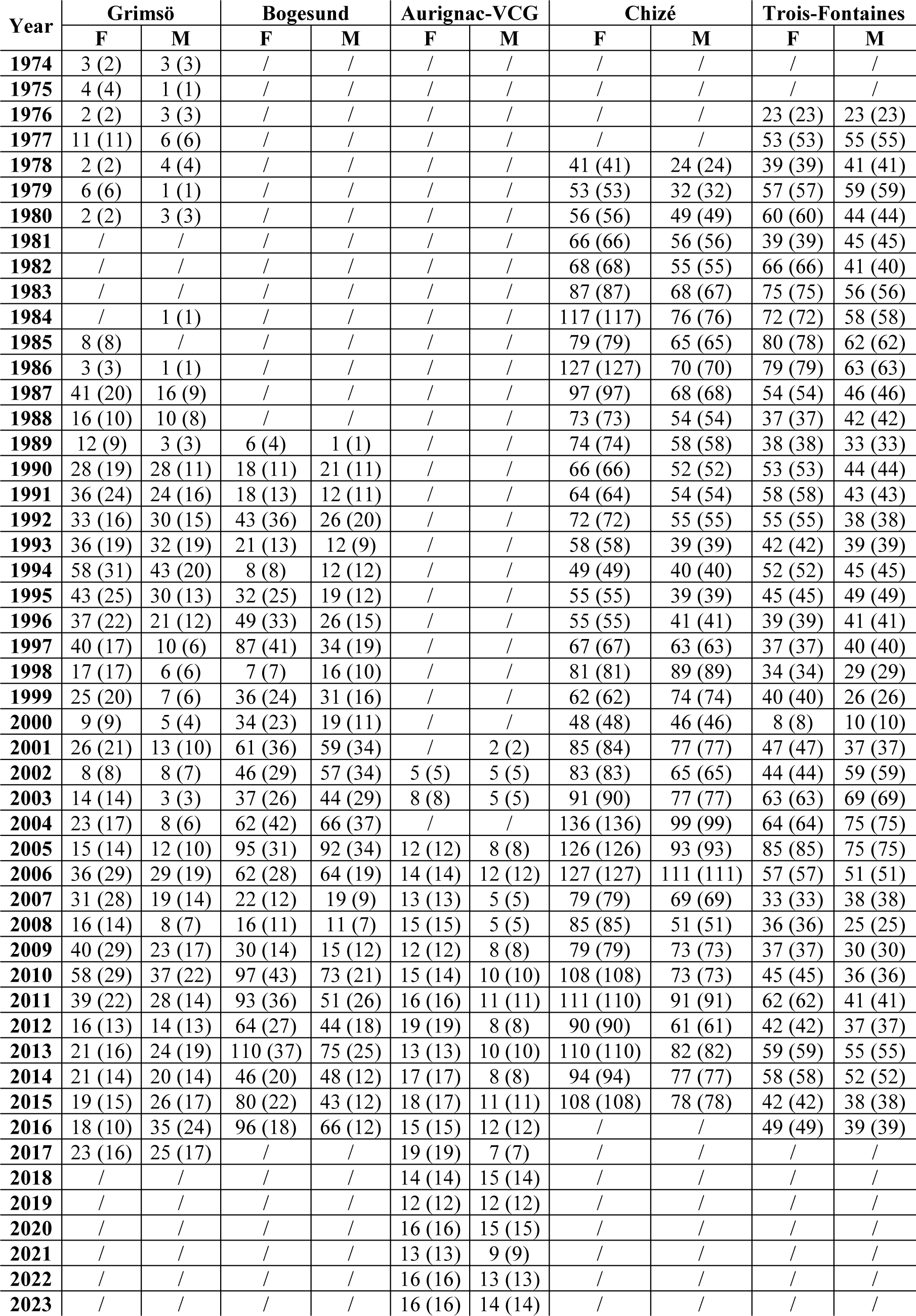
Sample size per year and sex for the five populations of roe deer situated in Sweden (Bogesund, Grimsö) and France (Aurignac-VCG, Chizé, Trois-Fontaines), with the number of unique individuals measured that year in brackets.

## Notes

https://doi.org/10.5281/zenodo.10997219

### Competing Interest Statement

The authors have declared no competing interest.

### Summary of Updates

This is the final version which has been recommended for publication. There are no major changes with respect to the previous version.

